# ABCC1 protects skin dendritic cells from FITC-induced toxicity by efflux and extracellular glutathione buffering

**DOI:** 10.64898/2026.01.02.697391

**Authors:** Konrad Knöpper, Anshul Rao, Jinping An, Jason G. Cyster

## Abstract

Dendritic cell (DC) migration is critical for initiating adaptive immune responses. Previous work suggested a role for ATP-binding cassette transporter C1 (ABCC1) in skin DC migration following cutaneous fluorescein isothiocyanate (FITC) exposure, but the precise mechanism involved was unclear. Here we establish that the primary contribution of ABCC1 to skin DC function following FITC exposure is not modulation of migration, but enhancement of survival. Our findings demonstrate that ABCC1 operates on a dual level: intracellularly, by transporting toxic FITC and fluorescein out of DCs, and extracellularly, by contributing to a glutathione (GSH) buffer zone that protects surrounding cells. DCs are particularly susceptible to FITC-mediated toxicity, possibly due to their high endocytic activity. This study elucidates the critical dependence of DCs on ABCC1 and extracellular GSH for resistance to toxic organic molecules and thereby identifies potential therapeutic avenues targeting ABCC1 to modulate immune responses.

**Significance Statement:** Adaptive immunity is critically dependent on dendritic cells (DCs) and their ability to take up foreign molecules for processing and presentation to T cells. DCs in the skin are exposed to a diversity of environmental chemicals and whether they have mechanisms to protect themselves from chemical-induced toxicity has been unclear. In this work we investigate the role of the multi-drug resistance transporter ABCC1 (MRP1) in DC biology. We reveal that ABCC1 shields DCs from chemical poisoning following skin exposure to fluorescein isothiocyanate (FITC). ABCC1 is needed both cell intrinsically and cell extrinsically to achieve the full protective effect. This discovery underscores a previously unrecognized reliance of DCs on ABCC1 function and opens new opportunities for therapeutic manipulation of these cells.

## Introduction

DCs in the skin continually sample their microenvironment for antigens and for the presence of pathogen associated molecular patterns (PAMPs), damage associated molecular patterns (DAMPs) and inflammatory cytokines. Upon exposure to such stimuli, they upregulate CCR7 and costimulatory molecules, migrate into skin lymphatics in a CCR7 dependent manner and subsequently into the T zone of the draining lymph node (LN) (Worbs et al., 2017). Here they present antigen-derived peptides on MHC molecules to prime T cell responses. The barrier location of DCs together with their endocytic activity and their need for metabolically intense behaviors may make them particularly sensitive to environmental chemicals.

ATP-binding cassette transporter C1 (ABCC1, also known as multi-drug resistance transporter protein-1 or MRP1) has a well-defined role in effluxing a range of anionic molecules out of cells, including various drugs and environmental chemicals, often as glutathione (GSH) conjugates (Cole, 2014). ABCC1 also effluxes the oxidized form of GSH, GSSG, and this helps reduce potentially toxic intracellular buildup of this metabolite. Additionally, ABCC1 contributes to the export of GSH into the extracellular milieu (Cole, 2014). Another important substrate of ABCC1 is the GSH conjugated bioactive lipid leukotriene C4 (LTC4) (Cole, 2014). In recent work, we established ABCC1 is a transporter of S-geranylgeranyl-L-glutathione (GGG), a ligand for P2RY8, in lymphoid tissues (Gallman et al., 2021). Whether ABCC1 has a role in exporting environmental chemicals from DCs and thereby protecting their immune function is unknown.

The fluorescein coupled form of isothiocyanate (FITC) has been used for over four decades to study skin DC activation and migration to skin draining LNs (sdLNs) (Macatonia et al., 1986; Thomas et al., 1980). Isothiocyanates (ITCs) are chemically reactive and can couple to thiol and amino groups (Hoch et al., 2024; Karlsson et al., 2016). Following topical application, various types of skin-derived DCs that harbor FITC-labeled proteins begin appearing in the sdLN within several hours and continue to accumulate over 2-3 days (Macatonia et al., 1987). The mechanism by which FITC induces DC maturation is not well understood but given its chemical reactivity it may involve triggering of DAMP sensing pathways.

Here we pursue the basis for the finding that ABCC1 is required for skin DC migration to LNs following exposure to FITC (Randolph et al., 2005; Robbiani et al., 2000; van de Ven et al., 2009). ABCC1 is highly expressed by DCs, and we found that the transporter protects DCs from toxicity and cell death induced by FITC and fluorescein. DCs were more sensitive than other CD45+ cell types to this toxicity and more dependent on ABCC1 for survival. Bone marrow (BM) chimera studies provided evidence that ABCC1 contributes both intrinsically and extrinsically to DC resistance to FITC-induced toxicity. These findings indicate that DCs have a high dependence on ABCC1-mediated molecular efflux and ABCC1-dependent extracellular GSH to sustain their viability and function following tissue exposure to some classes of reactive chemicals.

## Results

Prior work found that migratory DC accumulation in sdLNs 18 hours following FITC application on the skin was reduced in ABCC1-deficient mice (Robbiani et al., 2000). Using a similar FITC treatment approach we confirmed this observation in a distinct line of ABCC1-deficient mice (Fig. 1A and S1A). This reduction in migratory DCs was not evident at homeostasis and there was also no change in resident DCs or Langerhans cells (LCs) (Fig. 1B and S1B-E). The reduction in sdLN migratory DCs after FITC application was predominantly attributable to a diminished appearance of FITC+ migratory DCs (Fig. 1C and S1F). At this 18 hour time point the migratory skin DCs are conventional DCs as LCs have a delayed migratory behavior and take several days to reach the sdLN (Kissenpfennig et al., 2005; Shklovskaya et al., 2008). While a defect in ABCC1-deficient skin DC migration was previously suggested to involve impaired responsiveness to the CCR7 ligand CCL19 (Robbiani et al., 2000), other work found that CCL19 was dispensable for DC migration following FITC-mediated activation (Britschgi et al., 2010). Consistent with the latter findings, we did not observe a significant difference in the appearance of FITC+ DCs in the sdLN when comparing CCL19-deficient and control mice (Fig. 1D). Thus, the ABCC1-dependent DC phenotype appeared unlikely to be due to altered CCL19-directed migration.

**Figure 1.**
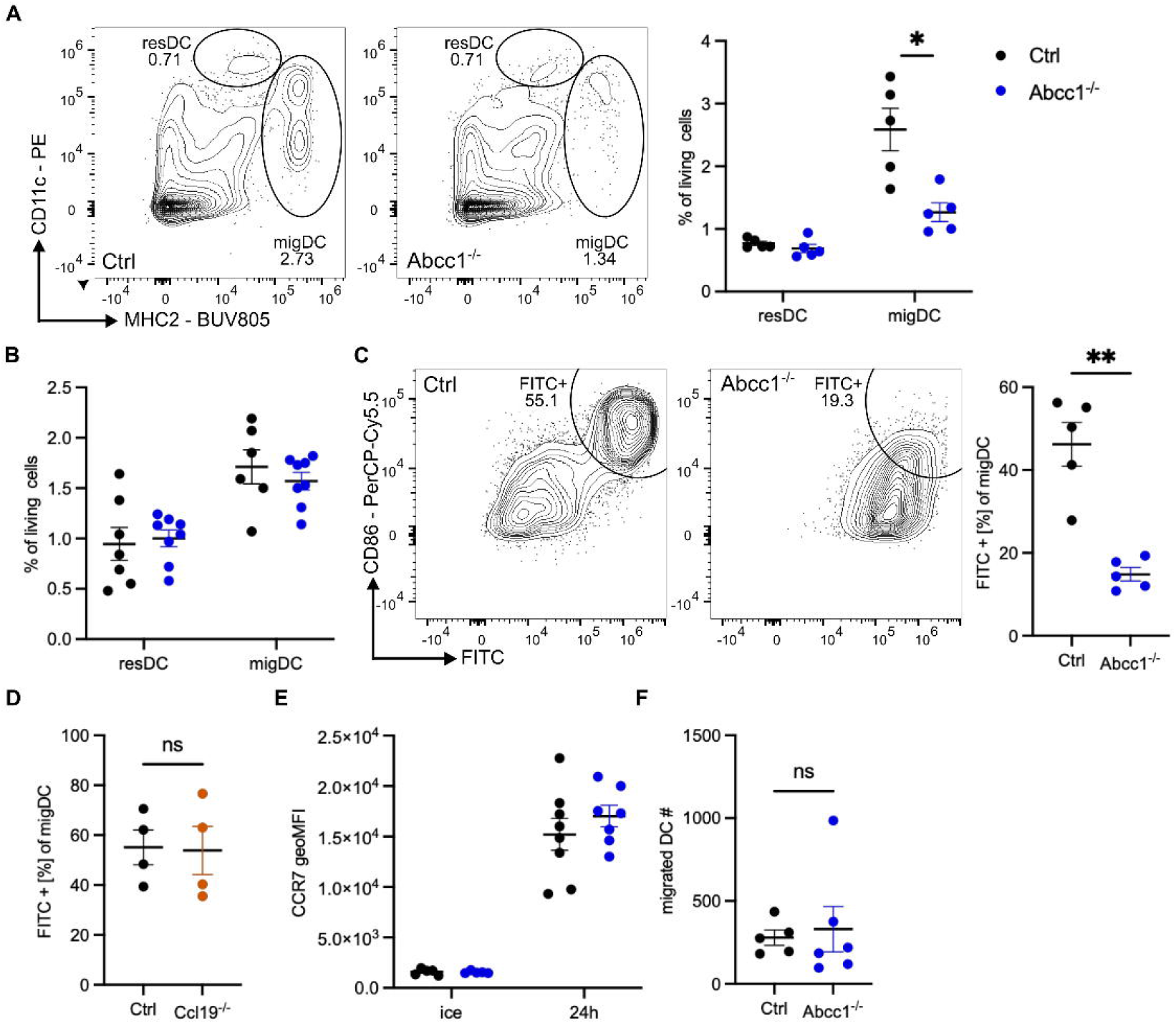
ABCC1 promotes accumulation of migratory DCs in the sdLN following FITC treatment in a CCL19 and migration independent manner. A) Representative flow cytometry plot and quantification of sdLN DC subsets in control and ABCC1-deficient mice. sdLN were analyzed 18 h after FITC treatment on the skin. B) Quantification of sdLN resident and migratory DC subsets in control and ABCC1-deficient naive mice. C) Representative flow cytometry plot and quantification of sdLN FITC+ migratory DCs in control and ABCC1-deficient mice. sdLN were analyzed 18 h after FITC treatment on the skin. D) Quantification of sdLN FITC+ migratory DCs in control and CCL19-deficient mice. sdLN were analyzed 18 h after FITC treatment on the skin. E) Quantification of CCR7 geoMFI staining from spleen DCs. DCs were isolated and either matured for 24 h in culture media or kept on ice. F) Quantification of migrated DCs from ear-crawl-out assay after 24 h. All data are representative of at least two independent experiments (n ≧ 3 mice per experiment). Error bars indicate SEM, and statistical analysis was done with Student’s t-test. ns = not significant ;*p-value < 0.05; **p-value < 0.01.

In accordance with this observation, we detected comparable surface CCR7 expression on *ex vivo* matured spleen DCs from ABCC1-deficient and control mice (Fig. 1E). Furthermore, analysis of DC migration via crawl-out assay from ear skin explants revealed no significant differences between ABCC1-deficient and control mouse skin (Fig. 1F). These observations collectively led us to consider that the migration of DCs may not be directly compromised by ABCC1-deficiency, and that the reduction in FITC+ DCs in sdLN reflected other roles for the transporter.

We next tested if alterations in DC behavior following FITC treatment might manifest within the skin. ABCC1-deficient mice harbored normal numbers of skin DCs at homeostasis (Fig. 2A) but showed a markedly reduced recovery of skin DCs 18 hours post-FITC application (Fig. 2B). The reduction in skin DC numbers in ABCC1-deficient mice was detectable as early as 4 hours post-treatment, arguing against the reduction being secondary to increased egress of DCs from the tissue (Fig. 2C). Skin LCs were also reduced 4 hours after FITC treatment (Figs. S2A and S2B). Moreover, activated DCs (CD86+) were almost entirely absent in FITC treated ABCC1-deficient mouse skin (Fig. 2D). This observation could not be reconciled by a defect in the activation capacity, as *ex vivo* maturation (Fig. S2C) and activation following exposure to lipopolysaccharide (LPS) and papain occurred comparably for ABCC1-deficient and control cells (Fig. 2E).

**Figure 2.**
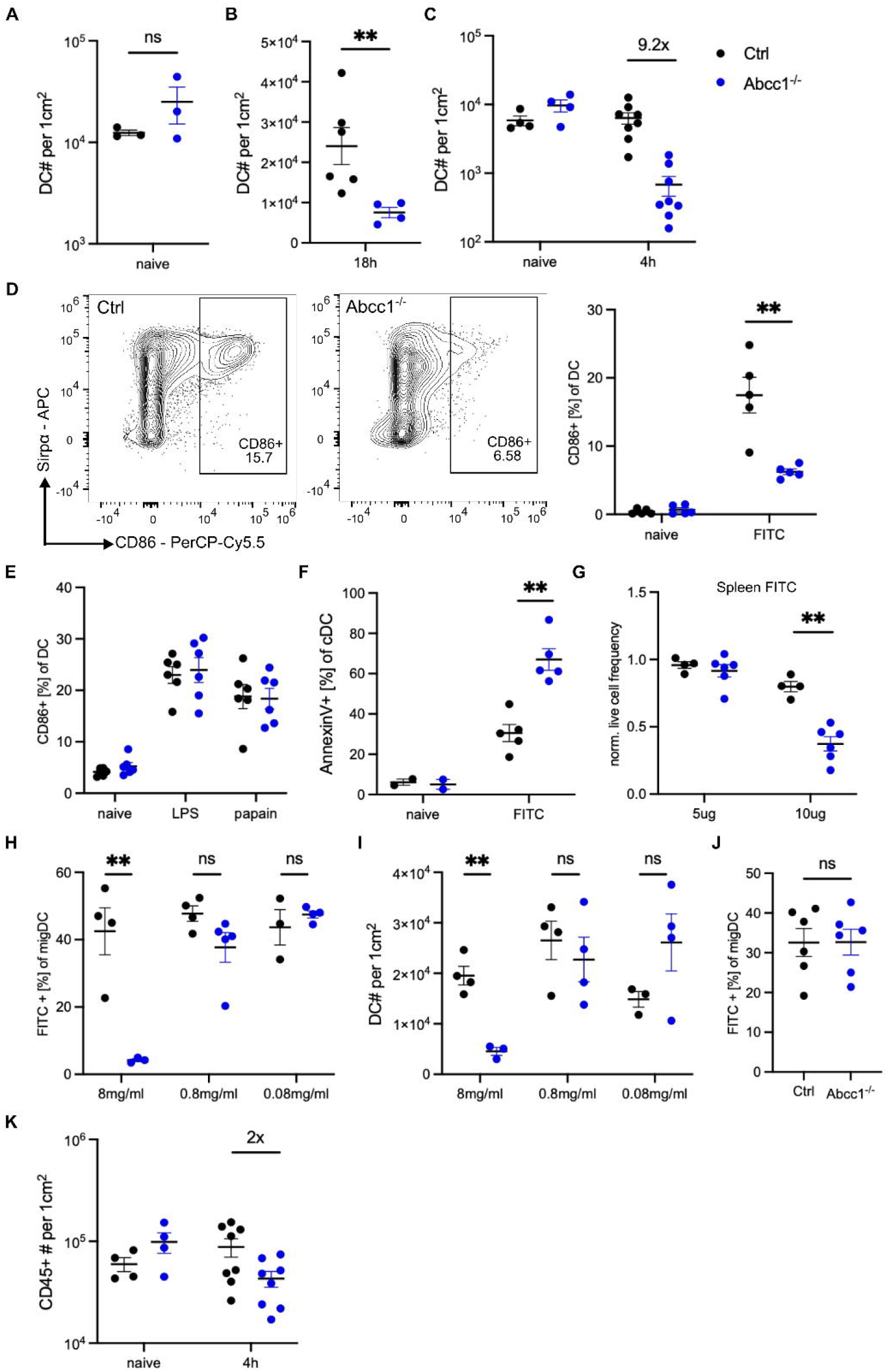
FITC treatment induces apoptosis in ABCC1-deficient DCs. A) Quantification of skin DCs in control and ABCC1-deficient naive mice. B) Quantification of skin DCs in control and ABCC1-deficient mice 18 h after FITC treatment. C) Quantification of skin DCs in control and ABCC1-deficient mice 4 h after FITC treatment. D) Representative flow cytometry plot and quantification of CD86 expression on skin DCs in control and ABCC1-deficient mice. Skin was analyzed 4 h after FITC treatment on the skin. E) Quantification of CD86 expression on skin DCs in control and ABCC1-deficient mice 4 h after papain or LPS treatment. F) Quantification of Annexin-V staining on skin DCs in control and ABCC1-deficient mice 4 h after FITC treatment. G)Quantification of normalized live cell frequency from splenocytes of control and ABCC1-deficient mice 24 h after FITC treatment *ex vivo*. Live frequency is normalized to solvent control. H)Quantification of sdLN FITC+ migratory DCs in control and ABCC1-deficient mice. sdLN were analyzed 18 h after skin treatment with the indicated FITC concentrations. I) Quantification of skin DCs in control and ABCC1-deficient mice 18 h after FITC treatment as in I. J) Quantification of sdLN FITC+ migratory DCs in control and ABCC1-deficient mice. sdLN were analyzed 3 days after 0.08 mg/ml FITC treatment on the skin. K) Quantification of skin CD45+ cells in control and ABCC1-deficient mice 18 h after FITC treatment. All data are representative of at least two independent experiments (n ≧ 3 mice per experiment). Error bars indicate SEM, and statistical analysis was done with Student’s t-test. ns = not significant; **p-value < 0.01.

We observed markedly greater accumulation of FITC in several cell types, following both *in vivo* and *ex vivo* FITC exposure, in ABCC1-deficient compared to control mice (Fig. S2D and S2E).

Given the elevated intracellular accumulation of FITC and the concomitant loss of DCs from the skin, we speculated that the treatment might have induced apoptosis in the DCs. Indeed, we detected increased Annexin-V surface staining of ABCC1-deficient DCs following FITC treatment (Fig. 2F). To assess whether this phenomenon could be attributed to secondary effects, we cultured splenocytes from control and ABCC1-deficient mice in the presence of FITC. Consistent with our *in vivo* data, ABCC1-deficient splenocytes exhibited reduced viability compared to control cells following incubation with FITC (Fig. 2G). We next compared the dose sensitivity of the FITC-induced effects on skin DCs. 10-fold or 100-fold lower amounts of FITC were sufficient to promote control skin DC migration (Fig. 2H). At these lower FITC doses ABCC1-deficiency did not impact DC migration and skin DC numbers were not reduced by the treatment (Fig. 2I). To exclude the possibility that FITC dilution merely delayed cell death, we measured the frequency of FITC+ DCs three days post-treatment with the reduced dose of FITC. No significant difference was observed in FITC+ DC frequency between ABCC1-deficient and control mice (Fig. 2J).

Importantly, the reduction in skin DC numbers was more pronounced than that observed for other cell types. While DCs were reduced 9.2-fold four hours post-FITC treatment in ABCC1-deficient mice (Fig. 2C), the total CD45+ cell population decreased by only 2-fold (Fig. 2K). To further investigate differential sensitivity to FITC-induced cell death, we compared DCs with dendritic epidermal T cells (DETCs), cells that are expected to be highly exposed to FITC due to their epidermal localization. Treatment of mice with 2 mg/mL FITC revealed a significant reduction in DC numbers in ABCC1-deficient mice, whereas DETC numbers remained comparable to control mice (Fig. 3A). These results suggest that DCs are more susceptible than lymphocytes to FITC induced apoptosis.

**Figure 3.**
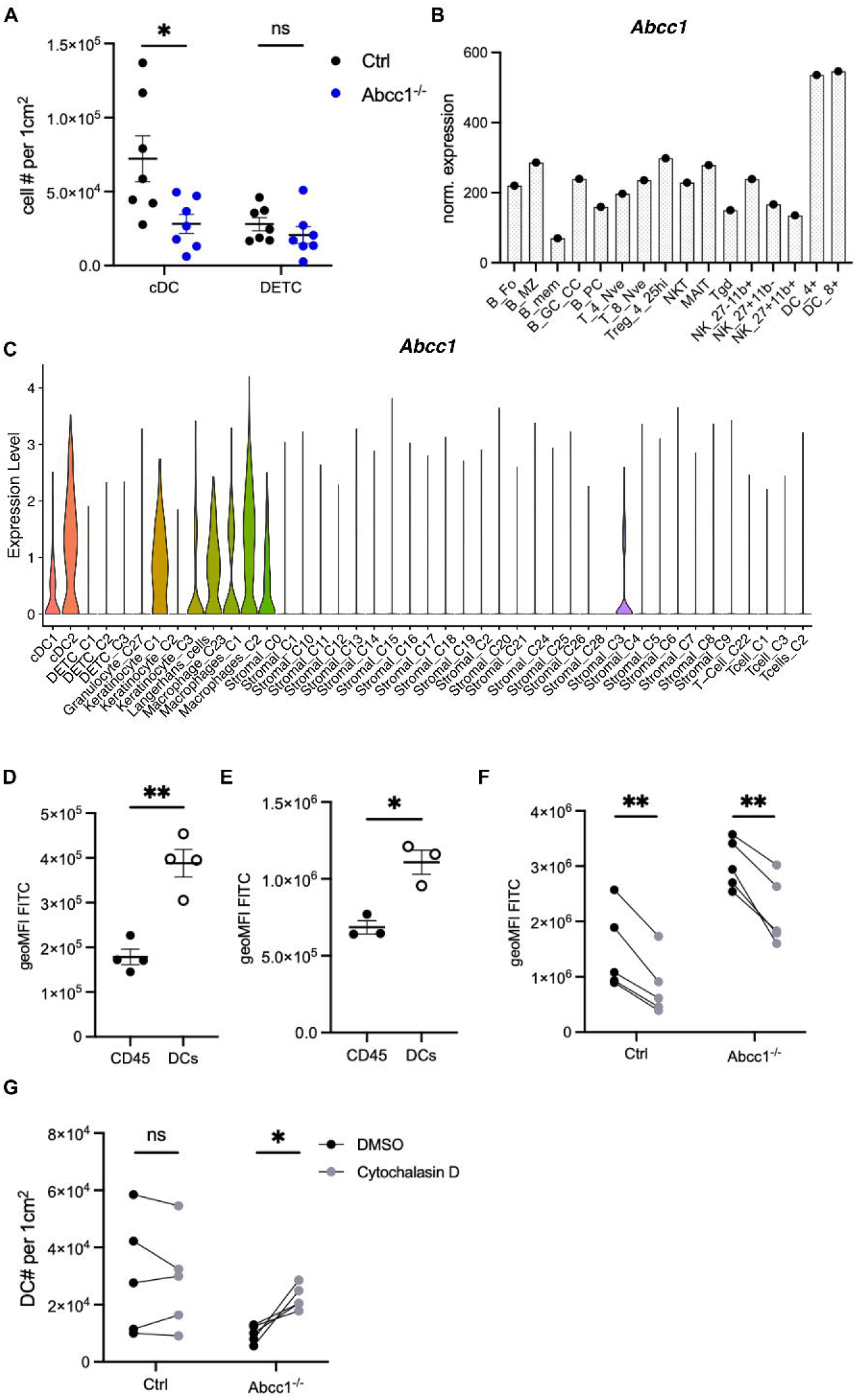
DCs have higher dependency on ABCC1 for survival. A) Quantification of skin DETC and DCs in control and ABCC1-deficient mice 4 h after FITC treatment. B) Normalized expression level of *Abcc1* in immune cells (immgen.org). C) Normalized expression level of *Abcc1* in skin single-cell data displayed as a violin plot. Cells are grouped by cell type. D) Quantification of FITC geoMFI of total CD45+ cells or DCs from spleen 24 h after FITC treatment *ex vivo*. E) Quantification of FITC geoMFI of total CD45+ cells or DCs from skin 4 h post FITC treatment. F) Quantification of FITC geoMFI of DCs from skin 4 h after FITC treatment. Groups were either treated with DMSO or Cytochalasin D. G) Quantification of skin DCs in control and ABCC1-deficient mice 4 h after FITC treatment. Groups were either treated with DMSO or Cytochalasin D. All data are representative of at least two independent experiments (n ≧ 3 mice per experiment). Error bars indicate SEM, and statistical analysis was done with Student’s t-test. ns = not significant; **p-value < 0.01; ***p-value < 0.001.

Analysis of publicly available spleen RNA-seq data (Immgen.org, (Heng et al., 2008)) indicated that DCs exhibited higher *Abcc1* expression than lymphocyte populations (Fig. 3B). Analysis of scRNA-seq data generated from mouse skin cells (Ben-Shaanan et al., 2024) revealed that skin DCs, LCs, and macrophages expressed *Abcc1* at higher levels than lymphoid cells or stromal cells (Fig. 3C and Fig. S3A, B). Collectively, these data suggest a greater requirement for ABCC1 function in DCs, LCs, and likely macrophages compared to other immune cells. In line with this notion, we observed that following FITC exposure *in vivo* and *ex vivo*, DCs harbored significantly higher FITC levels than the other immune cells that were analyzed (Fig. 3D-E).

Considering the differences in myeloid and lymphoid cell biology and the elevated intracellular FITC levels, we speculated that high endocytic activity (micropinocytosis), a hallmark of DCs, might underlie our observations. To investigate this possibility, we pharmacologically blocked micropinocytosis with the actin polymerization inhibitor Cytochalasin D (Qiu et al., 2022; Salloum et al., 2023). This treatment reduced FITC levels in both control and ABCC1-deficient DCs in the skin (Fig. 3F), but not significantly in DETCs (Fig. S3C), indicating micropinocytosis may be responsible for the higher FITC levels in DCs. Furthermore, inhibiting micropinocytosis partially rescued DCs from cell death, as evidenced by increased DC numbers in the skin of FITC-painted ABCC1-deficient mice (Fig. 3G). We do not exclude the possibility that Cytochalasin D treatment impacts additional cellular processes that influence DC survival.

Our data indicate that ABCC1 protects cells from apoptosis with the *in vitro* data providing evidence for a cell-intrinsic mechanism. To further test the cell intrinsic role of the transporter, we generated mixed chimeric mice by reconstituting lethally irradiated hosts with a 1:2 mixture of control or ABCC1-deficient BM cells and congenically marked wildtype BM cells. In the absence of challenge, the proportion of control or ABCC1-deficient skin DCs in the chimeras was comparable in skin (Fig. 4A) and sdLN (Fig. 4B). Following FITC challenge, there was a reduction in the proportion of ABCC1-deficient DCs in skin (Fig. 4C) and of ABCC1-deficient migratory DCs in sdLN (Fig. 4D). These data indicate that ABCC1 acts in a DC-intrinsic manner. However, the reduction in DCs was less prominent in the mixed chimera setting than in fully ABCC1-deficient mice (Fig. 4E) suggesting that ABCC1 also promotes DC survival in a cell-extrinsic manner.

**Figure 4.**
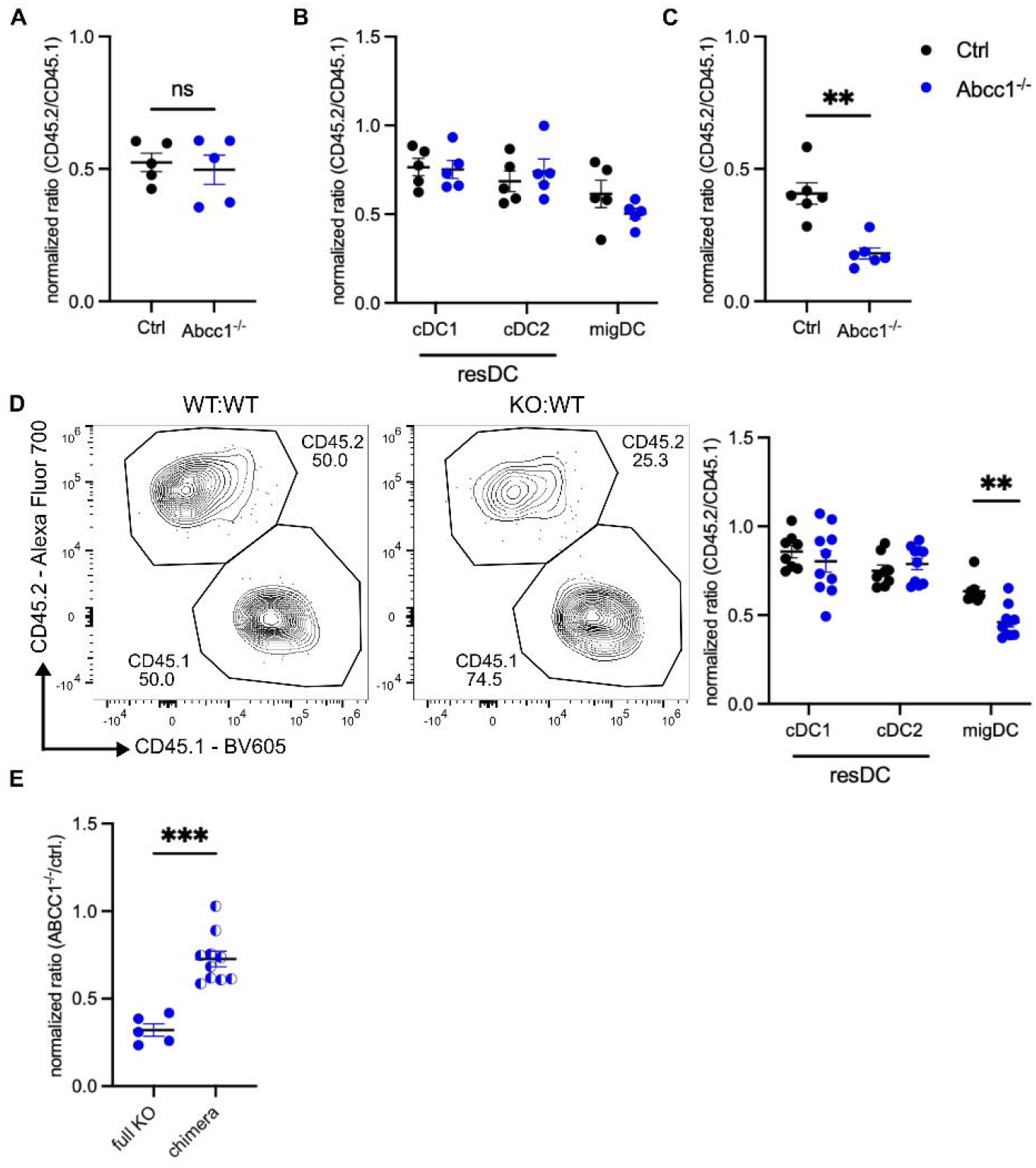
ABCC1 acts cell intrinsically and cell extrinsically to promote DC survival. (A, B) Quantification of normalized CD45.2/CD45.1 ratio of skin DCs (A) or sdLN DCs (B) in mixed BM chimeric mice with CD45.2 either representing control (Ctrl) or Abcc1^-/-^ cells and CD45.1 WT cells. (C, D) Quantification of normalized CD45.2/CD45.1 ratio as in A of skin DCs (C) and representative flow cytometry plot and quantification of normalized ratio of sdLN DCs (D) 18 h after FITC treatment. E) Normalized ratio of Abcc1^-/-^ to control sdLN DCs in FITC painted non-chimeric (full KO) and mixed chimeric mice. Data in E are from Fig. 1C and Fig. 4D. For non-chimeric mice the ratio is between different mice (Abcc1^-/-^ versus WT) whereas for mixed chimeric mice the ratio is for Abcc1^-/-^ versus WT cells in the same chimera. A-D) CD45.2/CD45.1 DC ratio is normalized to CD19+ cell ratio from the sdLN. All data are representative of at least two independent experiments (n ≧ 3 mice per experiment). Error bars indicate SEM, and statistical analysis was done with Student’s t-test. *p-value < 0.05; **p-value < 0.01.

One function of ABCC1 is the transport of GSH across the plasma membrane (Cole, 2014). Extracellular GSH contributes to the antioxidant properties of skin (Shindo et al., 1993). We speculated that extracellular GSH might play a role in protecting DCs from FITC-induced apoptosis by interacting with the ITC group of FITC and limiting its entry into the cells. To this end, we cultured splenocytes together with FITC and added GSH to the culture. Adding GSH reduced the FITC induced apoptosis (Fig. 5A). To test whether extracellular GSH has a protective effect *in vivo*, we subcutaneously treated control and ABCC1-deficient mice with GSH immediately before FITC painting. As before, we could recover less DCs from the skin in ABCC1-deficient mice 4 h post FITC painting compared to control mice (Fig. 5B). Treating with GSH ameliorated the phenotype and partially rescued the skin DCs from apoptosis (Fig. 5B). To further test the activity of ABCC1 in protecting DC viability cell-extrinsically we reconstituted ABCC1-deficient and control mice with a mixture of ABCC1-deficient and control BM-cells. This set-up allowed us to investigate the survival of ABCC1-deficient cells comparing a skin environment almost completely lacking GSH exported by ABCC1 to physiological extracellular GSH levels. FITC painting revealed that the ABCC1-deficient DC survival in the skin was reduced in a GSH deprived environment compared to control mice with an intact ABCC1 GSH export capacity (Fig. 5C and S4A). Together, these data provide evidence for the importance of extracellular GSH exported by ABCC1 in protecting DCs from FITC induced cell death.

**Figure 5.**
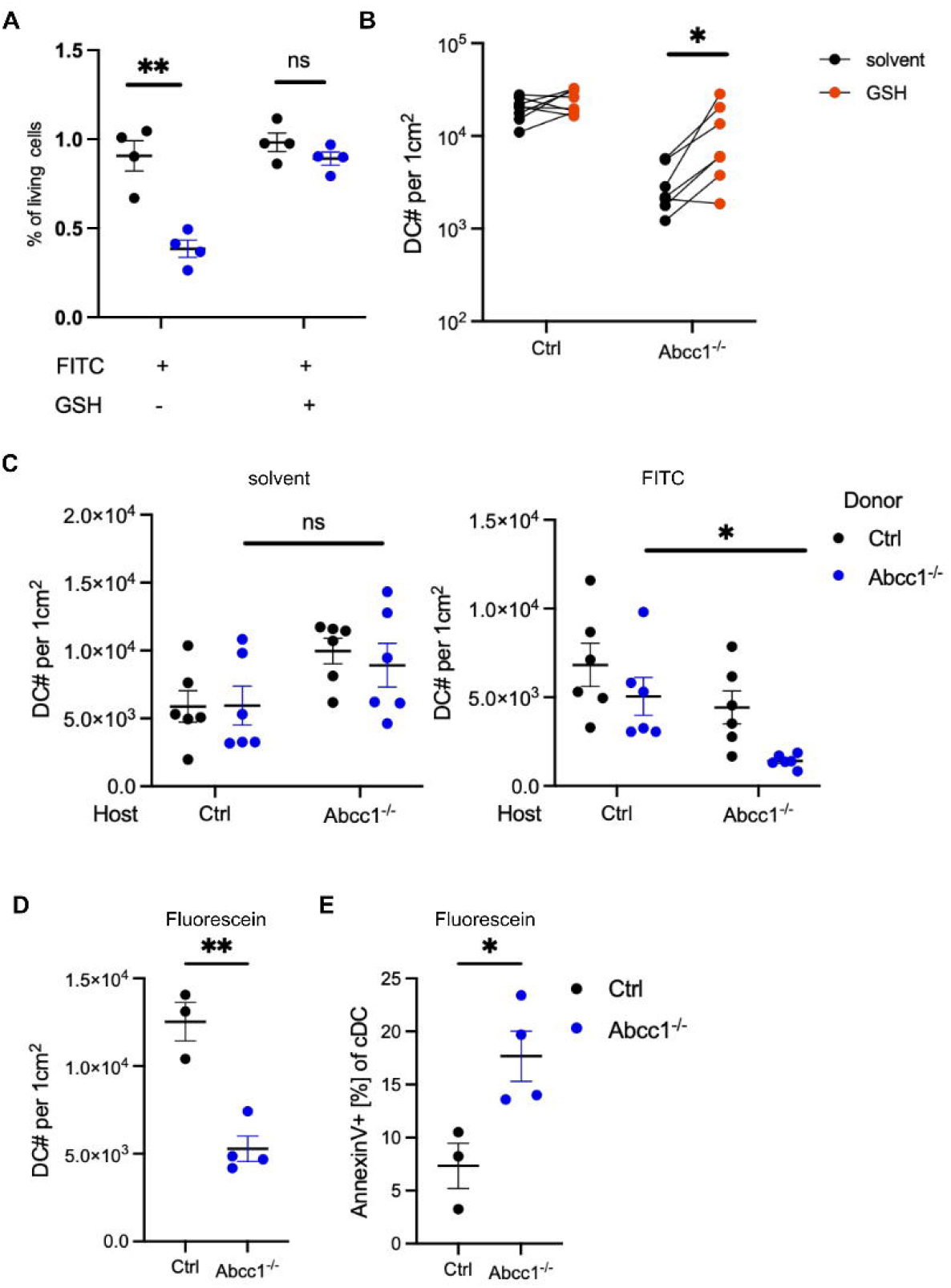
Activity of extracellular GSH in protecting DCs from FITC and sensitivity of ABCC1-deficient DCs to fluorescein. A) Quantification of normalized live cell frequency from splenocytes of control and ABCC1-deficient mice 18 h after FITC (10ug/ml) and GSH (50ug/ml) treatment *ex vivo*. B) Quantification of skin DCs in control and ABCC1-deficient mice 4 h after FITC and either saline or 1mg GSH treatment. C) Quantification of skin DCs with host mice either representing control or Abcc1^-/-^ and DCs being derived from control or Abcc1^-/-^ donor BM. Left plot depicts solvent treated mice and right depicts mice 4 h after FITC treatment. D) Quantification of skin DCs in control and ABCC1-deficient mice 4 h after fluorescein treatment. E) Quantification of Annexin-V staining on skin DCs in control and ABCC1-deficient mice 4 h after fluorescein treatment. All data are representative of at least two independent experiments (n ≧ 3 mice per experiment). Error bars indicate SEM, and statistical analysis was done with Student’s t-test. ns = not significant.

Finally, we tested the selectivity of ABCC1 in mediating DC protection from FITC-induced toxicity. ABCC1-deficiency did not affect DC sensitivity to benzylisothiocyanate (BITC) (Fig. S4B) or several other ITCs (PEITC, AITC, unpubl. obs.). To investigate whether FITC’s toxicity in ABCC1 deficient cells was independent of its ITC group, we treated mice with fluorescein. Fluorescein had a similar propensity as FITC to induce skin DC cell loss (Fig. 5D) and Annexin-V staining showed that this occurred by apoptosis (Fig. 5E). DETC were minimally altered in fluorescein sensitivity by the loss of ABCC1 (Fig. S4C). These data suggest that ABCC1 protects DCs from FITC toxicity at least in part by preventing toxic intracellular accumulation of fluorescein, independent of FITC’s ITC group. We then tested skin DC sensitivity to two chemotherapeutic agents that can be substrates for ABCC1, doxorubicin and vincristine (Cole, 2014). We did not observe increased sensitivity of ABCC1-deficient DCs to cell death induced by these chemicals. (Fig. S4D-E). These data indicate ABCC1 has a role in protecting skin DCs from a select class reactive chemicals.

## Discussion

Together our findings highlight the importance of ABCC1, also known as multi-drug resistance transporter protein-1 (MRP1), in enabling skin DCs to resist the toxic effects of FITC and fluorescein. Furthermore, we provide evidence for the extracellular function of GSH in reducing FITC reactivity and thereby promoting skin DC viability.

In earlier work that identified a role for ABCC1 in the migratory response of skin DCs following FITC exposure, a model was proposed involving ABCC1-exported LTC4 enabling CCR7-dependent migration to CCL19 (Robbiani et al., 2000). However, the CCL19 findings were based on *ex vivo* data and DC migratory responses are challenging to study *ex vivo* due to spontaneous DC maturation (Steinman et al., 2003). Analysis of CCL19-deficient mice does not support a requirement for CCL19 during FITC-induced skin DC migration *in vivo* ((Britschgi et al., 2010) and this report). Moreover, we found that skin exposure to low amounts of FITC induced equal migration of ABCC1-deficient and control DCs. A study tracing lung DC migration to the dLN following fluorescent bead exposure also showed no difference comparing ABCC1-deficient and control mice (Jakubzick et al., 2006). These findings favor the general conclusion that ABCC1 is not directly required for CCR7-mediated DC migration from tissue to dLNs.

Immature DCs undergo micropinocytosis and are estimated to take up 1000um^3^ or 40% of their volume every hour (Sallusto et al., 1995). By internalizing more interstitial fluid than other cell types DCs may experience increased entry of environmental chemicals to the cytosol where they could react with essential intracellular proteins, thereby making DCs more sensitive to chemical-induced toxicity. The high sensitivity of ABCC1-deficient DCs to FITC and fluorescein may also reflect their strong metabolic needs as they undergo maturation and migration within hours of exposure to these agents. Highly metabolically active cells may be more dependent on the intactness of their intracellular proteins and thus may experience stresses more readily than cells that are in a quiescent state.

While FITC is known to react with and label proteins in the plasma membrane, it can also enter cells (Edidin et al., 1976). In this regard, it is notable that FITC has a MW below 500 Da and thus within a size range that permits passage cross membranes (Matsson and Kihlberg, 2017; Wang et al., 2025). This is in accord with our finding that deficiency in ABCC1, a transporter that effluxes molecules out of the cytosol, causes cells to accumulate higher amounts of FITC. An open question remaining is how FITC or fluorescein are cytotoxic. Future studies will need to address the molecular mechanism and whether a DC-specific biological pathway is disrupted by these chemicals.

Glutathione (GSH), the most abundant thiol containing molecule in cells reacts with ITCs and modulates their toxicity (Hoch et al., 2024). GSH is also abundant extracellularly in skin (Shindo et al., 1993). ABCC1 contributes to extracellular GSH (Cole, 2014) and by reacting with FITC this may limit its access to cells. Our mixed chimera and *in vitro* culture data establish a cell intrinsic role for ABCC1 in limiting FITC accumulation in DCs and other cells, in accord with a role in efflux of FITC that reached the cytosol and likely reacted with GSH. However, our data also indicate a role for extracellular GSH in restraining FITC entry into cells, thus providing evidence that ABCC1 helps establish a chemical buffer zone in the skin. While GSH is the most likely ABCC1 substrate protecting cells from FITC, ABCC1 can transport a variety of other substrates, and we do not exclude the possibility that it influences the extracellular milieu in additional ways that protect cells from reactive chemicals.

By removing an important exporter of diverse cargo, we show that DCs have greater sensitivity than other cell types tested to one type of environmental chemical. The lack of ABCC1-deficient DC sensitivity to several other chemicals we tested may reflect other members of the ABC transporter family compensating for the loss of ABCC1. Future experiments will need to address the contribution of multiple ABC transporter genes to DC physiology and resistance to environmentally induced toxicity. Humans are exposed to a huge diversity of environmental chemicals including many new synthetic molecules. It may prove valuable for advancing understanding of the potential impact on these molecules on the immune system to screen them broadly for negative effects on control and ABC transporter-deficient DC viability.

## Materials and Methods

### Animals

Breeding was done in an internal colony, housed under 12-hour light: 12-hour dark cycles and given ad libitum access to food and water. 7 to 18-week-old mice of both sexes were used for experiments. Abcc1^-/-^ mice and Ccl19^-/-^ mice were generated as described previously and were on a C57BL6/J (B6) background (Link et al., 2007; Liu et al., 2022). B6 mice and CD45.1 congenic B6 mice (B6.SJL-PtprcaPepcb/BoyCrCrl) were bred internally. Littermate and cage mate controls were used for experiments. Animals were housed in a pathogen-free environment in the Laboratory Animal Resource Center at the University of California, San Francisco, and all experiments conformed to ethical principles and guidelines that were approved by the Institutional Animal Care and Use Committee.

### Treatment of mice

Mice were shaved with a clipper at the dorsal back skin before treatment. GSH (5mg), Cytochalasin D (1ug), papain (50ug), and LPS (5ug) were injected subcutaneously in 50ul at the stated concentrations dissolved in saline or saline with 1% DMSO. FITC, fluorescein, BITC, vincristine and doxorubicin were applied in 25ul to the shaved dorsal skin at stated concentrations. For topical applications, the chemicals were dissolved in a 1:1 mix of Dibutylpthalate:Acetone and freshly prepared before treatment.

### BM-chimera generation

Mice, where indicated, were lethally irradiated with 900 rads gamma-irradiation (split dose separated by 4 h) and then i.v. injected with 5x10^6 BM cells. To generate mixed BM-chimera, BM cells from each donor mouse were counted using a hemocytometer and 5x10^6 BM-cells were transferred in total with the mixing ratio indicated in the main text. Mice were analyzed 6-8 weeks post reconstitution.

### Ear-crawl out assay

Ears were harvested from mice and split to separate ventral and dorsal half. Each half was placed, dermis facing down, onto 1 ml of RPMI with 1mM Hepes, and 10% fetal bovine serum, supplemented with P/S, GlutMax and NEAA in a 6-well plate. The tissue was incubated for 24 h and media was collected. Dendritic cell numbers in media were counted using Cytek aurora after staining with flow antibodies.

### Cell isolation LN

Mice were sacrificed, LNs were isolated and placed in 1ml digestion buffer (composed of 10 ug/ml collagenase IV and 0.1mg/ml DNase in RPMI with 1mM Hepes, and 2% fetal bovine serum). The LNs were rotated at 37°C for 60 minutes. Afterwards, a single cell suspension was generated using a pestle and the cells were transferred into staining buffer (PBS containing 2% NBCS and 0.05% Sodium Azide) for cell counting and staining.

### Cell isolation skin

A shaved skin patch of 1cmx1cm was dissected, minced finely with scissors, re-suspended in 1 ml of digestion media (composed of 0.25mg/ml Liberase TM, 0.5mg/ml hyaluronidase and 0.1mg/ml DNase in RPMI with 1mM Hepes, and 2% fetal bovine serum), and incubated in a heated shaker at 37°C at 1000 rpm for 60 minutes. An additional 1ml of RPMI/Hepes/FBS media containing 1mM EDTA was added and the suspension, vortexed and then strained into a 50ml conical tube through a 70um strainer. Another 10ml of media was added to thoroughly wash the cell strainer and the suspension was centrifuged, re-suspended in staining buffer (PBS containing 2% NBCS and 0.05% Sodium Azide) for cell counting and staining. Cell counts per 1cm^2^ of digested skin were derived from Cytek aurora analyzer and calculated based on acquired volume.

### Splenocyte isolation and culture

Spleens were harvested, minced finely with scissors, re-suspended in 1 ml of digestion media (composed of 10 ug/ml collagenase IV and 0.1mg/ml DNase in RPMI with 1mM Hepes, and 2% fetal bovine serum). The spleens were rotated at 37°C for 60 minutes. Afterwards, a single cell suspension was generated using a pestle. Red blood cells were lysed using RBC lysis buffer and afterwards the single cell suspension was transferred into staining buffer (PBS containing 2% NBCS and 0.05% Sodium Azide). Where indicated, splenocytes were cultured in culture media (composed of RPMI with 1mM Hepes, and 10% fetal bovine serum, supplemented with P/S, GlutMax and NEAA).

### Flow cytometry analysis

Single cell suspension obtained from tissues were used for flow cytometry analysis using Cytek aurora. The data was analyzed and visualized using Flowjo software (v10). Cells were surface stained with Zombi NIR stain (Biolegend), anti-γδTCR (Biolegend; clone: GL3; BV650), anti-CD11b (BD; clone: M1/70; BUV493), anti-CD11c (Biolegend; clone: N418; PE/ Spark NIR 685/ APC), anti-CD19 (BD; clone: 1D3; BUV563), anti-IA/IE (BD; clone: M5/114; BUV805/ PerCP-cy5.5), anti-Epcam (Biolegend; clone: G8.8; BV711), anti-CD64 (Biolegend; clone: W18349C; PE-Cy7), anti-CD86 (Biolegend; clone GL-1; PerCP-Cy5.5), anti-Sirpa (Biolegend; clone: P84; PE-Dazzle594/ APC), anti-XCR1 (Biolegend; clone: ZET; AL647), anti-CD45.2 (Biolegend; clone: 104; BUV395/ AL700), anti-CD45.1 (Biolegend; clone: A20; AL700/ BV605), Ly6G (Biolegend; clone: 1A8; APC-Cy7), CD4 (BD; clone: GK1.5; BUV615), CD8a (BD; clone: 53-6.7; BUV737).

### CCR7 stain

Anti-CCR7 antibody (Biolegend; clone: 4B12; biotinylated) was incubated with cells for 1 h at 37°C. Afterwards, cells were washed with PBS containing 2% NBCS and 0.05% Sodium Azide two times. SA conjugated to PE (Biolegend) was incubated together with additional surface antibodies in staining buffer for 30 minutes on ice.

### Annexin-V staining

After surface antibody staining, cells were washed with Annexin-V staining buffer (BD) and stained in Annexin-V staining buffer with PE-conjugated Annexin-V (BD) for 15 minutes at RT. Cells were washed with Annexin-V staining buffer and analyzed using Cytek aurora.

### RNA-sequencing data processing and analysis

For the presented bulk and scRNA-sequencing results, published datasets were analyzed running a customized R script with Seurat v.4.2 for single cell sequencing. Raw counts of the published data were used (GEO Accession: GSE266028, GSE266027) or normalized expression values were retrieved from immgen.org.

### Statistical analysis

Apart of the genomic data, all biological data were analyzed using Prism 10 software (GraphPad) by two-tailed paired Student’s t-test or two-tailed unpaired Student’s t-test.

## Acknowledgements

We thank current and former Cyster lab members for helpful discussion and feedback. KK is supported by a DFG Walter Benjamin postdoctoral fellowship. JGC is an Investigator of the Howard Hughes Medical Institute. This work was supported in part by NIH grant AI 40098.

## References

Ben-Shaanan, T.L., K. Knopper, L. Duan, R. Liu, H. Taglinao, Y. Xu, J. An, M.V. Plikus, and J.G. Cyster. 2024. Dermal TRPV1 innervations engage a macrophage- and fibroblast-containing pathway to activate hair growth in mice. Dev. Cell 59:2818–2833 e2817.

Britschgi, M.R., S. Favre, and S.A. Luther. 2010. CCL21 is sufficient to mediate DC migration, maturation and function in the absence of CCL19. Eur. J. Immunol. 40:1266–1271.

Cole, S.P. 2014. Targeting multidrug resistance protein 1 (MRP1, ABCC1): past, present, and future. Annu. Rev. Pharmacol. Toxicol. 54:95–117.

Edidin, M., Y. Zagyansky, and T.J. Lardner. 1976. Measurement of membrane protein lateral diffusion in single cells. Science 191:466–468.

Gallman, A.E., F.D. Wolfreys, D.N. Nguyen, M. Sandy, Y. Xu, J. An, Z. Li, A. Marson, E. Lu, and J.G. Cyster. 2021. Abcc1 and Ggt5 support lymphocyte guidance through export and catabolism of S-geranylgeranyl-L-glutathione. Science Immunology 6:eabg1101.

Heng, T.S., M.W. Painter, and C. Immunological Genome Project. 2008. The Immunological Genome Project: networks of gene expression in immune cells. Nat. Immunol. 9:1091–1094.

Jakubzick, C., F. Tacke, J. Llodra, N. van Rooijen, and G.J. Randolph. 2006. Modulation of dendritic cell trafficking to and from the airways. J. Immunol. 176:3578–3584.

Kissenpfennig, A., S. Henri, B. Dubois, C. Laplace-Builhe, P. Perrin, N. Romani, C.H. Tripp, P. Douillard, L. Leserman, D. Kaiserlian, S. Saeland, J. Davoust, and B. Malissen. 2005. Dynamics and function of Langerhans cells in vivo: dermal dendritic cells colonize lymph node areas distinct from slower migrating Langerhans cells. Immunity 22:643–654.

Link, A., T.K. Vogt, S. Favre, M.R. Britschgi, H. Acha-Orbea, B. Hinz, J.G. Cyster, and S.A. Luther. 2007. Fibroblastic reticular cells in lymph nodes regulate the homeostasis of naive T cells. Nat. Immunol. 8:1255–1265.

Liu, D., L. Duan, L.B. Rodda, E. Lu, Y. Xu, J. An, L. Qiu, F. Liu, M.R. Looney, Z. Yang, C.D. Allen, Z. Li, A. Marson, and J.G. Cyster. 2022. CD97 promotes spleen dendritic cell positioning and homeostasis through sensing of red blood cells. Science 375:eabi5965.

Macatonia, S.E., A.J. Edwards, and S.C. Knight. 1986. Dendritic cells and the initiation of contact sensitivity to fluorescein isothiocyanate. Immunology 59:509–514.

Macatonia, S.E., S.C. Knight, A.J. Edwards, S. Griffiths, and P. Fryer. 1987. Localization of antigen on lymph node dendritic cells after exposure to the contact sensitizer fluorescein isothiocyanate. Functional and morphological studies. J. Exp. Med. 166:1654–1667.

Matsson, P., and J. Kihlberg. 2017. How Big Is Too Big for Cell Permeability? J. Med. Chem. 60:1662–1664.

Qiu, Z., W. Liu, Q. Zhu, K. Ke, Q. Zhu, W. Jin, S. Yu, Z. Yang, L. Li, X. Sun, S. Ren, Y. Liu, Z. Zhu, J. Zeng, X. Huang, Y. Huang, L. Wei, M. Ma, J. Lu, X. Chen, Y. Mou, T. Xie, and X. Sui. 2022. The Role and Therapeutic Potential of Macropinocytosis in Cancer. Front Pharmacol 13:919819.

Randolph, G.J., G. Sanchez-Schmitz, and V. Angeli. 2005. Factors and signals that govern the migration of dendritic cells via lymphatics: recent advances. Springer Semin. Immunopathol. 26:273–287.

Robbiani, D.F., R.A. Finch, D. Jager, W.A. Muller, A.C. Sartorelli, and G.J. Randolph. 2000. The leukotriene C(4) transporter MRP1 regulates CCL19 (MIP-3beta, ELC)-dependent mobilization of dendritic cells to lymph nodes. Cell 103:757–768.

Salloum, G., A.R. Bresnick, and J.M. Backer. 2023. Macropinocytosis: mechanisms and regulation. Biochem. J. 480:335–362.

Sallusto, F., M. Cella, C. Danieli, and A. Lanzavecchia. 1995. Dendritic cells use macropinocytosis and the mannose receptor to concentrate macromolecules in the major histocompatibility complex class II compartment: downregulation by cytokines and bacterial products. J. Exp. Med. 182:389–400.

Shindo, Y., E. Witt, and L. Packer. 1993. Antioxidant defense mechanisms in murine epidermis and dermis and their responses to ultraviolet light. J. Invest. Dermatol. 100:260–265.

Shklovskaya, E., B. Roediger, and B. Fazekas de St Groth. 2008. Epidermal and dermal dendritic cells display differential activation and migratory behavior while sharing the ability to stimulate CD4+ T cell proliferation in vivo. J. Immunol. 181:418–430.

Steinman, R.M., D. Hawiger, and M.C. Nussenzweig. 2003. Tolerogenic dendritic cells. Annu. Rev. Immunol. 21:685–711.

Thomas, W.R., A.J. Edwards, M.C. Watkins, and G.L. Asherson. 1980. Distribution of immunogenic cells after painting with the contact sensitizers fluorescein isothiocyanate and oxazolone. Different sensitizers form immunogenic complexes with different cell populations. Immunology 39:21–27.

van de Ven, R., G.L. Scheffer, R.J. Scheper, and T.D. de Gruijl. 2009. The ABC of dendritic cell development and function. Trends Immunol. 30:421–429.

Wang, Z., B.S. Pan, R.K. Manne, J. Chen, D. Lv, M. Wang, P. Tran, T. Weldemichael, W. Yan, H. Zhou, G.M. Martinez, J. Shao, C.C. Hsu, R. Hromas, D. Zhou, Z. Qin, H.K. Lin, and H.Y. Li. 2025. CD36-mediated endocytosis of proteolysis-targeting chimeras. Cell 188:3219–3237 e3218.

Worbs, T., S.I. Hammerschmidt, and R. Forster. 2017. Dendritic cell migration in health and disease. Nat. Rev. Immunol. 17:30–48.

