## Supplementary material for "ABCC1 protects skin dendritic cells from FITC-induced toxicity by efflux and extracellular glutathione buffering": SI

**This PDF file includes:**

Figures S1 to S4

### Figures

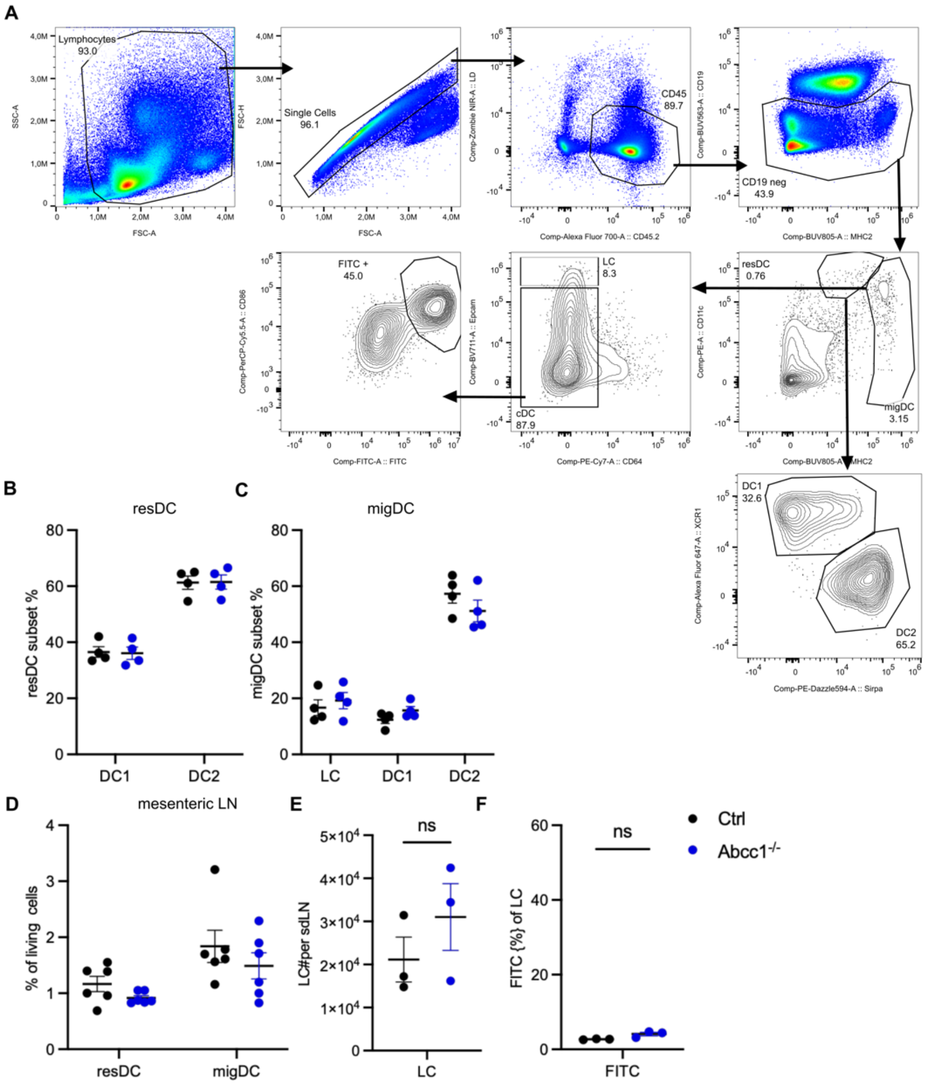


**Figure S1.** A) Gating strategy to identify FITC+ migratory DCs in the sdLN 18 h after FITC treatment. B) Quantification of sdLN resDCs subsets in control and ABCC1-deficient naive mice. C) Quantification of sdLN migDCs subsets in control and ABCC1-deficient naive mice. D)Quantification of mesLN DCs in control and ABCC1-deficient naive mice. E) Quantification of sdLN LC absolute numbers in control and ABCC1-deficient mice 18 h after FITC treatment. F) Quantification of sdLN FITC+ LC frequencies in control and ABCC1-deficient mice 18 h after FITC treatment. B-F is representative of one to two independent experiments (n ≧ 2 mice per experiment). Error bars indicate SEM.

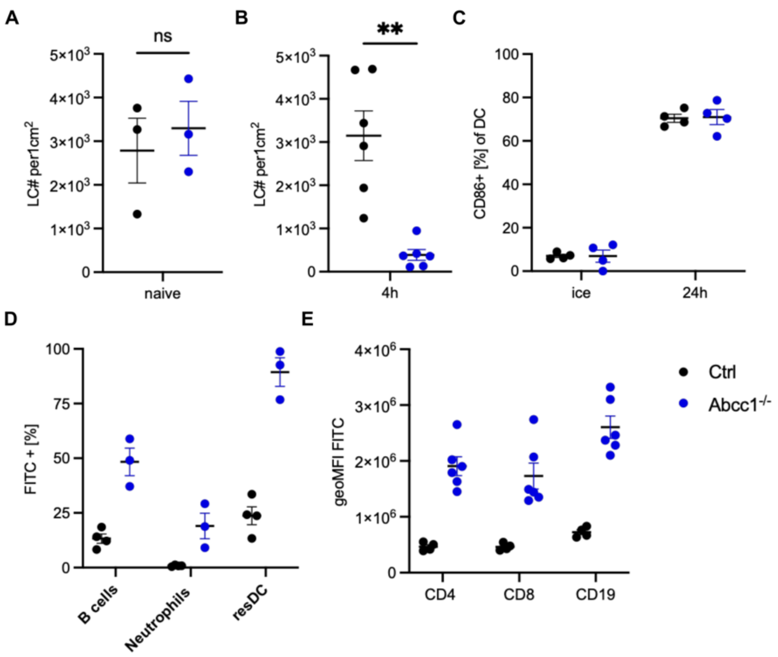


**Figure S2.** A) Quantification of skin LC in control and ABCC1-deficient naive mice. B) Quantification of skin LC in control and ABCC1-deficient mice 4 h after FITC treatment. C) Quantification of CD86 expression on spleen DCs in control and ABCC1-deficient mice after 24 h ex vivo culture. D) Quantification of FITC+ frequencies of sdLN cells 18 h after FITC treatment. E) Quantification of FITC geoMFI of splenocytes 24 h after FITC treatment ex vivo. All data are representative of at least two independent experiments (n ≧ 3 mice per experiment). Error bars indicate SEM.

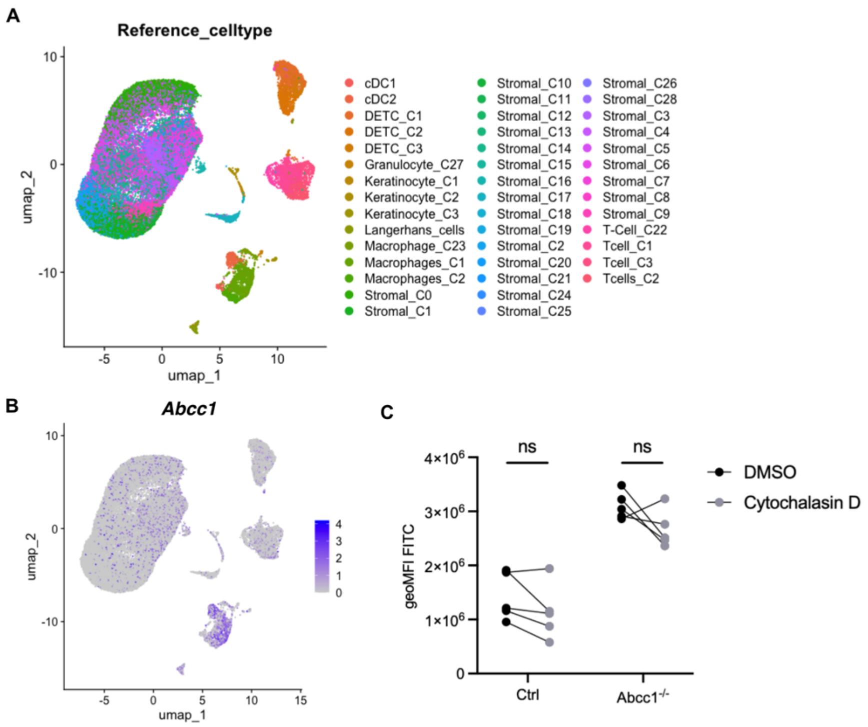


**Figure S3.** A) UMAP display of scRNA-sequencing data. Each dot represents a cell, colored by cell identity. B) UMAP display of scRNA-sequencing data. Each dot represents a cell, colored by Abcc1 expression level. C) Quantification of FITC geoMFI of DETC from skin 4 h after FITC treatment. Groups were either treated with DMSO or Cytochalasin D. All data are representative of at least two independent experiments (n ≧ 3 mice per experiment). Statistical analysis was done with Student's t-test. ns = not significant.

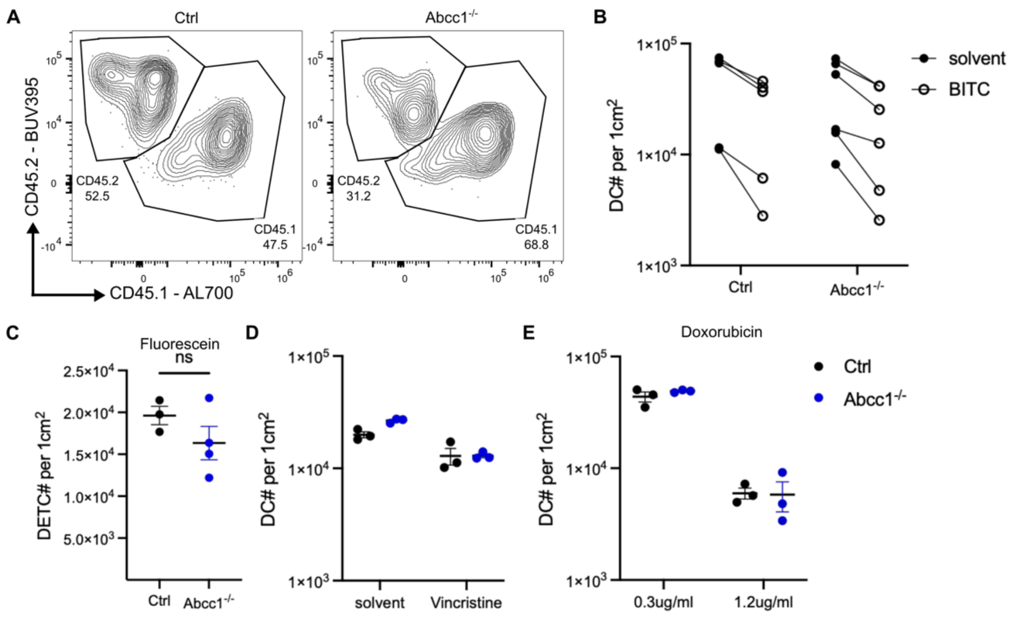


**Figure S4.** A) Representative flow cytometry plots showing CD45.1 (Ctrl) and CD45.2 (Abcc1-/-) ratio of skin DCs either from Ctrl (left) or ABCC1-deficient (right) hosts 4h after FITC treatment. B) Quantification of skin DCs in control and ABCC1-deficient mice 4 h after 2 % BITC treatment. C) Quantification of skin DETC in control and ABCC1-deficient mice 4 h after fluorescein treatment. D, E) Quantification of skin DCs in control and ABCC1-deficient mice 4 h after vincristine (20ug/ml) (D) or doxorubicin (E) treatment. All data are representative of at least two independent experiments (n ≧ 3 mice per experiment). Error bars indicate SEM, and statistical analysis was done with student t-test.
